# Optogenetic control shows that kinetic proofreading regulates the activity of the T cell receptor

**DOI:** 10.1101/432740

**Authors:** O. Sascha Yousefi, Matthias Günther, Maximilian Hörner, Julia Chalupsky, Maximilian Wess, Simon M. Brandl, Robert W. Smith, Christian Fleck, Tim Kunkel, Matias D. Zurbriggen, Thomas Höfer, Wilfried Weber, Wolfgang W.A. Schamel

## Abstract

The pivotal task of the immune system is to distinguish between self and foreign antigens. The kinetic proofreading model (KPR) proposes that T cells discriminate self from foreign ligands by the different ligand binding half-lives to the T cell receptor (TCR). It is challenging to test KPR as the available experimental systems fall short of only altering the binding half-lives and keeping other parameters of the ligand-TCR interaction unchanged. We engineered an optogenetic system using the plant photoreceptor phytochrome B to selectively control the dynamics of ligand binding to the TCR by light. Combining experiments with mathematical modeling we find that the ligand-TCR interaction half-life is the decisive factor for activating downstream TCR signaling, substantiating the KPR hypothesis.

**One Sentence Summary:** The half-life of the ligand-T cell receptor complex determines T cell activation.

## Main Text

The function of T cells is to mount an immune response to foreign ligands, such as derived from bacteria or viruses, but not to respond to self ligands stemming from the body’s own cells. Activation of a T cell is initiated when the foreign peptides presented by major histocompatibility complexes (MHC) on the own cells bind to the T cell receptor (TCR) on the T cell surface. This binding event stimulates intracellular signaling pathways leading to the functional responses of the T cell (*1*). Self peptides on MHC also bind to the TCR and are important for the survival of naïve T cells, but do not trigger an immune response as seen for foreign peptides (*2*). This discrimination between foreign and self peptides correlates with the affinity of the ligand-TCR interaction, in that foreign, stimulatory peptide-MHCs bind with higher affinity to the TCR than non-stimulatory peptide-MHC (*2, 3*). However, how the affinity of a ligand is determined by the cell to generate a T cell response or not remains enigmatic (*4*).

The kinetic proofreading (KPR) model originally described the specificity by which the genetic code is read in protein synthesis (*5*) and inspired a similar theoretical model for ligand discrimination in T cells (*6*). This model proposes that a long half-life of the ligand-TCR interaction, such as seen for high affinity peptides, allows a series of biochemical reactions to be completed that eventually trigger downstream signaling. By contrast, a low affinity ligand detaches before an activatory signal is produced and the TCR then reverts quickly to the initial inactive state, thus not initiating T cell activation. To get experimental insight into the mechanism of ligand discrimination by T cells, peptide-MHC or TCRs have been mutated at the binding sites to generate ligand-TCR pairs of different affinities and half-lives (*7–11*). Although such studies are broadly consistent with KPR, the free binding energy, geometry of the interaction (*12*), conformational changes at the TCR (*13*) and forces that act on the TCR (*14, 15*) might also have been changed along with the affinity, and therefore alternative models of ligand discrimination cannot be ruled out. In order to disentangle the half-life from these other parameters, we engineered an optogenetic system in which the duration of ligand binding to the TCR can be remotely controlled in a reversible manner (ON-OFF switch).

Our approach harnesses the PhyB-PIF (phytochrome B-PhyB interacting factor) protein pair from *Arabidopsis thaliana* (*16–18*). The PhyB-PIF interaction is turned on with 660 nm and turned off with 740 nm wavelength light, both within seconds (*16, 19, 20*). We fused PIF to the ectodomain of TCRb, as we did earlier with a single chain Fv fragment (*21*). Although soluble TCR ligands are active as dimers (*21, 22*), tetrameric peptide-MHC based on streptavidin are routinely used to stimulate the TCR (*23*). Similarly, streptavidin-based PhyB tetramers (PhyBt) act here to bind and cross-link the TCRs in a light-dependent manner (Fig. 1A and S1). While PIF is produced in the cytoplasm of plants, in our system it is present in the secretory pathway. Therefore, we needed extensive engineering of the PIF-TCRb construct to get a PhyBt-responsive TCR expressed on the surface of Jurkat T cells, referred to as GFP-PIF^S^-TCR (Supplementary Text and Fig. S2 to S4).

**Fig. 1.**
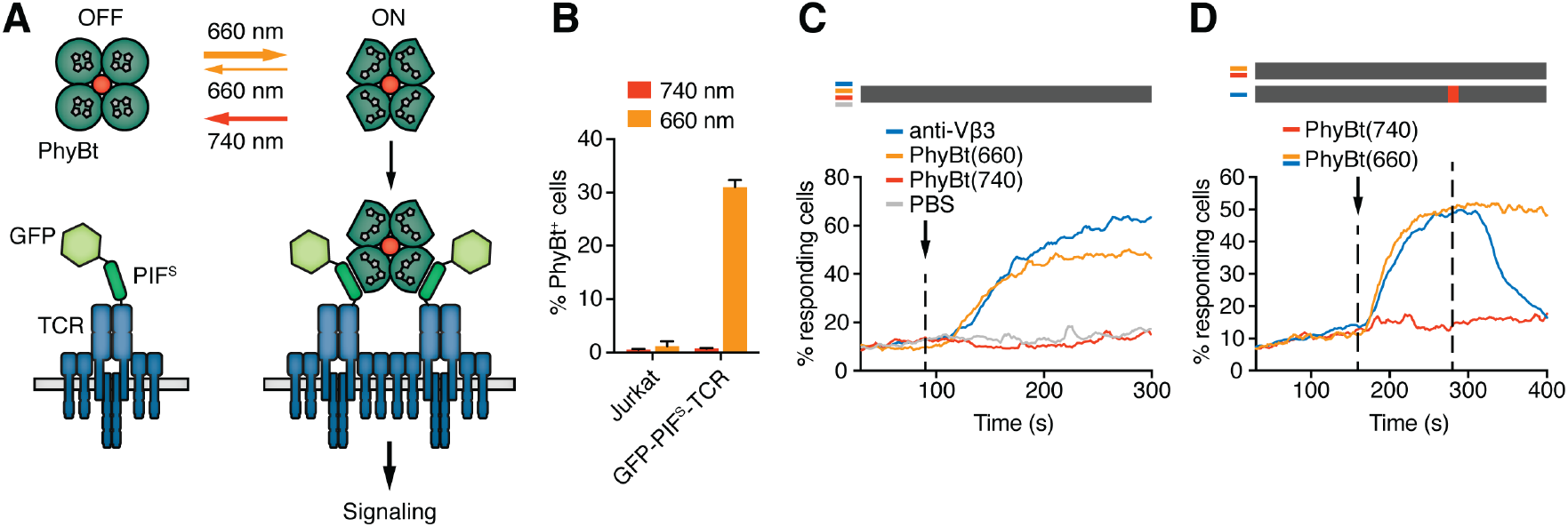
Engineering a light-controlled switch for the ligand-TCR interaction. (**A**) Light of 660 nm and 740 nm wavelength reversibly switches PhyB between the OFF and ON states. In the ON state PhyB tetramers (PhyBt) bind to and cluster GFPPIF^S^-TCRs leading to signaling and the activation of the T cell. (**B**) Fluorophore-labeled PhyBt pre-illuminated with 660 nm light [PhyBt(660)] bound to GFP-PIF^S^-TCR cells and PhyBt pre-illuminated with 740 nm light [PhyBt(740)] did not, as detected by flow cytometry. Jurkat cells serve as a control. The average of quadruplicates ± SEM is shown for one representative experiment of n>3. (**C**) Addition of PhyBt(660) (orange line) or anti-TCRVb3 antibody (blue line) to GFP-PIF^S^-TCR cells caused calcium influx. PhyBt(740) (red line) or PBS (grey line) did not induce a calcium response. The black arrow marks addition of the stimuli, and the bar above the graph represents the illumination procedure during the measurement (grey = dark). (**D**) PhyBt(660) added to GFP-PIF^S^-TCR cells induced calcium influx (blue and orange lines). After 2 min a 1 s short pulse of 100% intensity 740 nm light (red break in the grey bar) terminated the calcium response (blue line). Addition of PhyBt(740) did not induce calcium influx (red line). Results in (C) and (D) show one representative experiment of n>3.

PhyBt pre-illuminated with 660 nm wavelength light [PhyBt(660)] bound to cells expressing the GFP-PIF^S^-TCR, whereas PhyBt pre-illuminated with 740 nm light [PhyBt(740)] did not (Fig. 1B and S4F). Binding induced TCR signaling, since addition of PhyBt(660), but not PhyBt(740), resulted in a strong calcium influx into the cells similar to a stimulation using an anti-TCR antibody (Fig. 1C and S5A). The experiment was done in the dark, since in the absence of any light, the PhyB molecules rest in their state (ON or OFF) for time scales exceeding the duration of the calcium experiments (*19, 24*). 660 nm light alone in the absence of PhyBt or GFPPIF^S^-TCR did not evoke signaling, showing that the light acted through inducing PhyBt binding to GFP-PIF^S^-TCR (Fig. S5B and S5C). Further, stimulation with PhyBt under 660 nm light resulted in up-regulation of the activation marker CD69 (Fig. S5D). Together these data show that light-mediated PhyBt-binding to GFP-PIF^S^-TCR induced TCR signaling and T cell activation.

The PhyB-PIF system allows the rapid switching between the ON and OFF states in both directions. When we switched PhyBt from the ON to the OFF state by a 1 s pulse of 740 nm light, we stopped the ongoing calcium response initially evoked by PhyBt(660) (Fig. 1D), demonstrating that our system is reversible.

The KPR model predicts that the half-life of the ligand-TCR interaction determines TCR signaling. Here, we wanted to implement a protocol to control this half-life by light and study the consequences for TCR signaling, testing the KPR model. To this end, we exploited the property of PhyB that its continuous exposure to 660 nm light triggers both the switch from PhyB OFF to ON and the reverse switch from ON to OFF as the absorption spectra of both PhyB states partially overlap (*25*) (Fig. 2A). Thus, each individual PhyB molecule constantly shuttles between the ON and OFF state under 660 nm light, with high 660 nm intensities leading to a faster shuttling rate and thus to shorter binding duration (note that in Fig. 1 continuous light was not used and the PhyB ON molecules stayed in their state for longer times than relevant for the calcium experiments). Accordingly, continuous high intensity (100%) 660 nm light prevented calcium influx when PhyBt(660) was added to the GFP-PIF^S^-TCR cells (Fig. 2B, blue line). After 390 s the constant 660 nm illumination was stopped, so that the PhyB molecules that were in the ON state at this moment were trapped in this state. This allowed them to bind long enough to the TCR and to induce a strong calcium response (Fig. 2B). This experiment also demonstrates that the constant high intensity 660 nm illumination did not harm the cells.

**Fig. 2.**
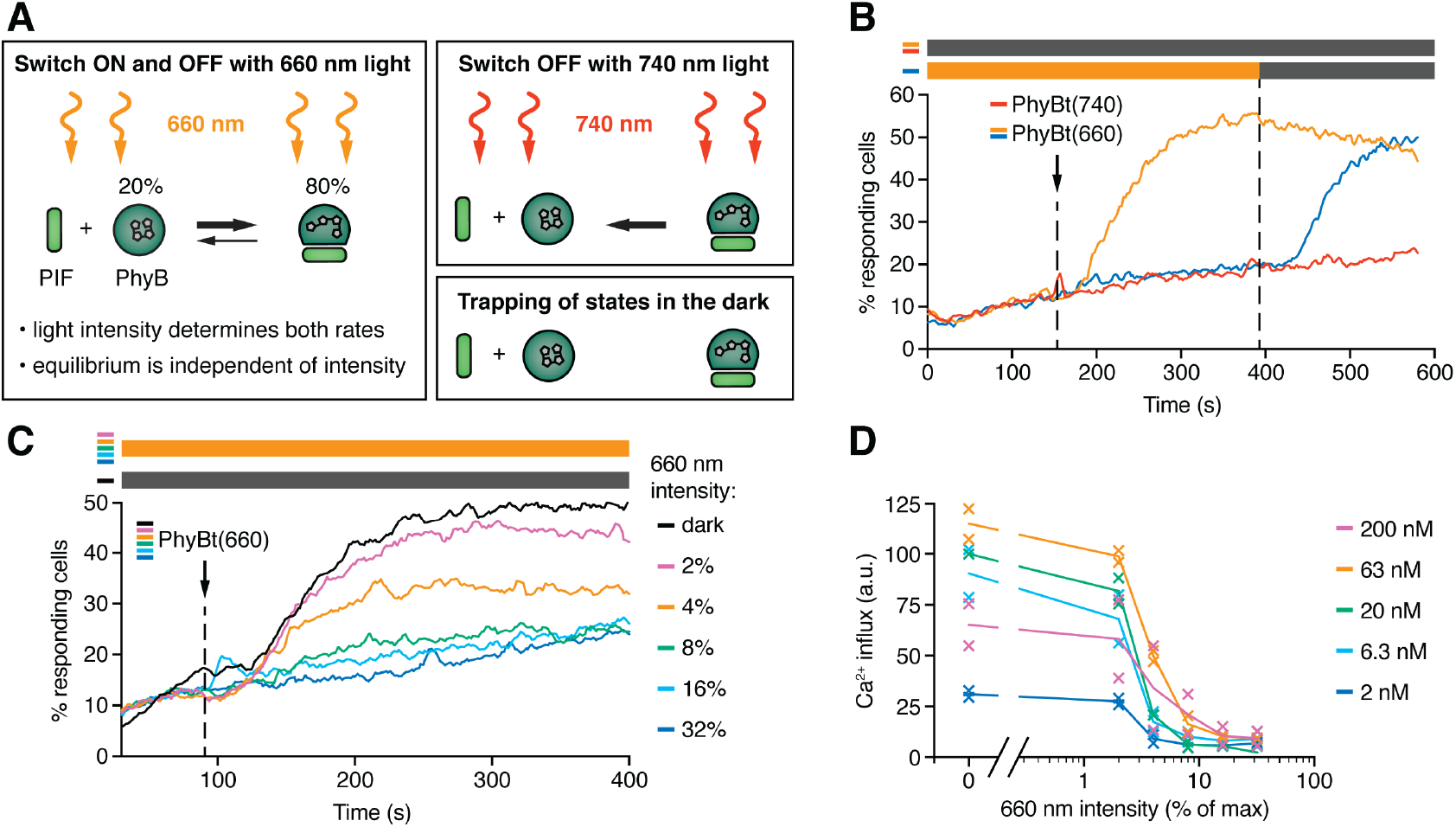
The half-life of the ON state of PhyB determines TCR signaling. (**A**) Schematics of the different PhyB conversions under 660 and 740 nm light. In the dark the PhyB states do not change in the timescales relevant for this work. (**B**) GFP-PIF^S^-TCR cells were constantly illuminated with 100% intensity 660 nm light (blue line). After 150 s PhyBt(660) was added (arrow) and after 390 s the light was switched off. As controls, PhyBt(660) (orange line) or PhyBt(740) (red line) was added to the cells in the dark. The bars represent the illumination procedure during the measurement (grey = dark, orange = 660 nm light). (**C**) 20 nM PhyBt(660) was added (arrow) after 90 s to GFP-PIF^S^-TCR cells continuously illuminated with 660 nm light of the depicted intensities. Results in (B) and (C) show one representative experiment of n>3. (**D**) Quantification of experiments done as in (C) with the indicated PhyBt concentrations. Duplicates are shown with connecting lines going through the mean.

The intensity of 660 light determines the half-life of both PhyB states and consequently the switch rates between the ON and the OFF state. However, the 80:20 molar ratio of PhyB ON to OFF molecules at photoequilibrium is largely independent of the light intensity (Fig. 2A) (*18, 19*). Lowering the 660 nm intensity increases the half-life of PhyB ON without altering its concentration, and hence may allow PhyBt to bind for longer durations to the GFP-PIF^S^-TCR. Indeed, at 4% and 2% constant 660 nm intensity, calcium influx was evoked (Fig. 2C). We observed a threshold of the PhyB ON half-life in inducing a calcium response that was largely independent of the PhyBt concentration, a crucial property of TCR ligand discrimination (*6*) (Fig. 2D and S6). In conclusion, we were able to control TCR signaling by changing the intensity of 660 nm light, suggesting that the duration of the ligand-TCR interaction controls calcium signaling.

Next, we developed a mathematical model and confronted it with the experimental data, to obtain quantitative insight into how the half-life of the PhyB ON-TCR complex determines TCR signaling. The model comprises the PhyB ONOFF cycle, binding of PhyBt ON to the TCR, and, potentially, KPR (Fig. 3A, S7, S8 and Supplementary Text). In the absence of KPR, the activity of each component in the signaling network depends only on the activity of its immediate upstream component(s), making TCR occupancy the ultimate source of ligand discrimination. In contrast, KPR assumes that the first signaling steps at the receptor in addition depend on the half-life of the ligand-TCR complex, while only the more downstream components respond exclusively to the activity of their immediate upstream component(s). We refer to the time required to complete the first half-life-dependent signaling steps as proofreading duration, τ_KPR_. Effectively, KPR requires the PhyB ONTCR complex to exist for at least the proofreading duration, in order to generate a signal that then leads to a calcium response more downstream (*6–8, 11*) (Fig. 3A). Thus, the time delay between ligand binding and calcium influx consists of the KPR duration plus the extra time beyond KPR required for the additional signaling steps until opening of the calcium channels. The half-life of the PhyB ON-TCR complex is determined by the sum of the light-independent off-rate of PhyB ON from the TCR, *k*_off_, and the light intensity-dependent rate *k*_i_ with which PhyB molecules return to the OFF state, detaching from the TCR (Fig. 3B).

**Fig. 3.**
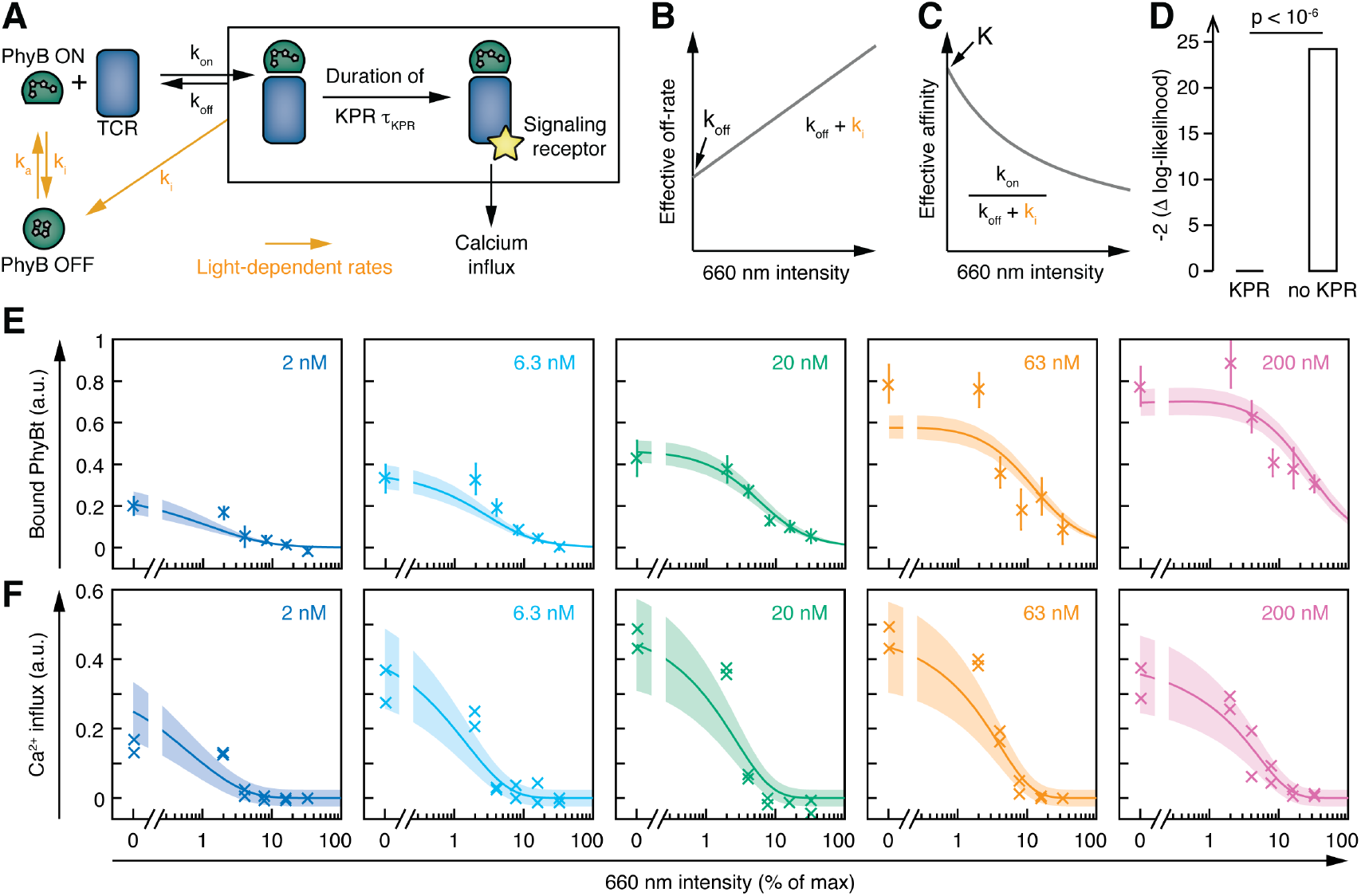
T cells exploit a kinetic proofreading mechanism. (**A**) The PhyB ON-OFF cycle, binding of PhyB ON to the TCR and kinetic proofreading (KPR) were combined into one model. (**B**) The effective off rate and **(C)** the effective affinity, both regulated by 660 nm light, are displayed as a function of the light intensity. (**D**) Comparing the best fit of the null hypothesis (no KPR, τ_*KPR*_ = 0) against the best fit of the alternative hypothesis (KPR, τ_*KPR*_ > 0) with a likelihood ratio test shows strong support for the existence of a KPR mechanism. (**E**), (**F**) Indicated concentrations of PhyBt(740) were added to GFP-PIF^S^-TCR cells and pre-illuminated with 660 nm light to fully activate PhyBt. The amount of PhyBt bound to the GFP-PIF^S^-TCR cells (**E**) and calcium influx (**F**) at different continuous 660 nm light intensities (from Fig. 2D) are plotted, however, in contrast to Fig. 2D, normalization is based on scaling factors estimated from the model. The line and shaded area represent the fit and the estimated uncertainties of the KPR model. The data points represent the mean ± SEM of triplicates in (**E**), n>5, or individual data points of two experiments in (**F**).

In support of the model (Fig. 3A), we experimentally demonstrated that the rate of converting PhyB from ON to OFF is the same for free PhyB and PIF-bound PhyB (Fig. S9). These data imply that PhyB molecules also convert to OFF while being bound to PIF and thereby the PhyB-PIF interaction is lost. Hence, the effective off-rate and binding affinity of PhyB ON to the TCR are also light-dependent (Fig. 3B and 3C). Taken together, the model predicts that the amount of TCR-bound PhyB decreases with increasing light intensity, which we confirmed experimentally (Fig. S10). Importantly, the PhyB ON affinity change is a straightforward consequence of the light-controlled PhyB ON half-life (this contrasts with mutated peptide-MHC ligands (*7–11*), where affinity changes can be brought about by changes in both on- and off-rates, and potentially other parameters such as orientation of binding (*12*)).

Although we intended to only change the ligand-TCR half-life with light, we also changed the affinity, due to the intrinsic relationship between off-rate and affinity. Hence, the intensity of 660 nm light regulates both the half-life of PhyB ON and the amount of bound PhyBt. To disentangle the half-life from the amount of ligand-bound TCRs, we asked whether calcium signaling was directly sensitive to the PhyB ON half-life through KPR or solely responded to the level of TCR occupancy with PhyB ON (absence of KPR). We fitted both mathematical models, the one with and the one without KPR, to the PhyBt binding and calcium signaling data together. Only the model with KPR yielded a satisfactory fit, and a likelihood ratio test between KPR and no-KPR models showed highly significant support for the KPR model (p < 10^−6^, Fig. 3D to 3F). The experimental data were fitted well when we included at least three KPR steps (Fig. S11, Supplementary Text). Since an upper bound for the number of KPR was not revealed by our combined experimental-modeling approach, we conclude that at least three fast KPR steps operate rather than very few longer lasting steps. Taken together, these findings strongly support the existence of KPR at the TCR.

The steady-state data (cf. Fig. 3E and 3F) prevented the model to deduce the proofreading duration τ_KPR_, yielding only the product τ_KPR_ *k*_off_. To overcome this limitation, we determined the conversion kinetics of PhyB in our experimental system by illuminating PhyBt OFF with short light pulses of 660 nm light and subsequently switching to darkness. This protocol traps the ligands in the ON state, which we quantified through the resulting calcium signal (Supplementary Text, Fig. S12A). The resulting kinetics of switching PhyB to the ON state were highly consistent across different light intensities (Fig. 4A) and PhyBt concentrations (Fig. S12B) and were described well by the mathematical model. Utilizing these data, we determined the half-life of the PhyB ON-TCR complex, ln2 /(*k*_off_ + *k*_i_), which varied from 40 s to 2 s over the range of light intensities used (Fig. 4B). The proofreading duration τ_KPR_ was then determined as 16 s [95% CI: 7 s, 38 s] (Fig. 4C). Thus, for a threshold half-life of the PhyB ON-TCR complex of 16 s, signaling from the active TCR is half-maximal. Furthermore, our results largely exclude the possibility of fast rebinding events, which would have effectively prolonged the half-life of the PhyB ON-TCR complex sensed by a KPR mechanism (*26, 27*) (Supplementary Text).

**Fig. 4.**
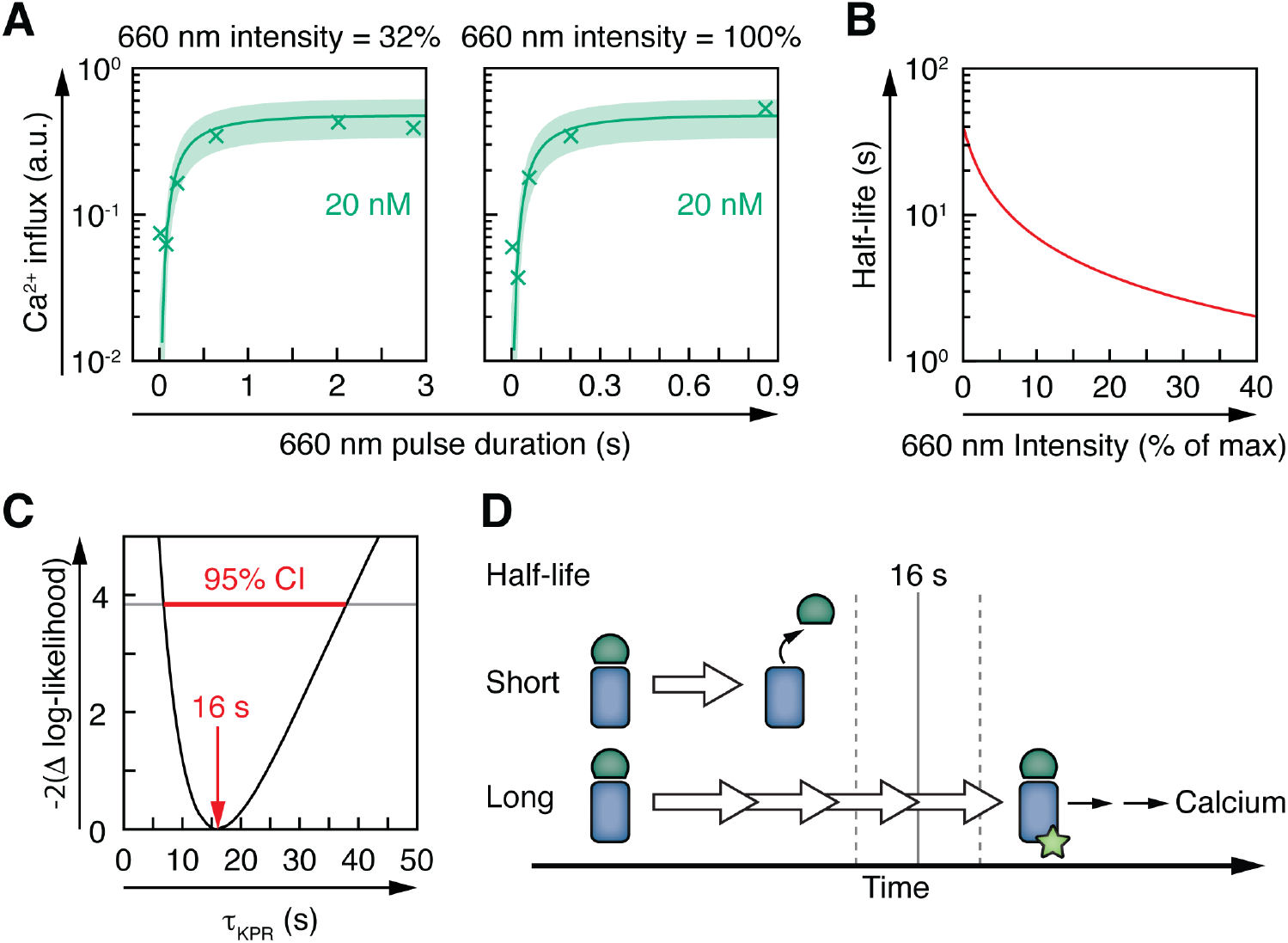
Kinetic proofreading at the TCR takes 16 seconds. (**A**) 20 nM PhyBt(740) was added to GFP-PIF^S^-TCR cells and pre-illuminated with the indicated pulse duration of 660 nm light at the indicated intensity settings to only partially activate PhyB. The induced calcium influx was measured in darkness, revealing the PhyBt conversion dynamics. The data is shown together with the fit and estimated SD. Results show one representative experiment of n>3. (**B**) Knowledge of the PhyBt conversion dynamics allows estimation of the half-lives of the PhyB-TCR complex in dependence on the light intensity. (**C**) The profile likelihood of the KPR duration shows that the 95% confidence interval (CI) ranges from 7 s to 38 s, while the best-fit value is 16 s. (**D**) Ligands that bind shorter than the KPR duration of 16 s fail to induce efficient TCR signaling. Ligands that bind longer allow the completion of at least three biochemical steps that result in T cell activation.

In this study, we engineer a tailor-made optogenetic system to control a ligand-receptor interaction by light, allowing us to overcome current experimental limitations. Our approach exploits the remarkable, but in optogenetics so far unexplored, biophysical property of PhyB that the intensity of 660 nm light determines the half-life of the PhyB ON state (*18, 19, 25*). Thereby, we controlled the ligand binding events in a non-synchronous manner mimicking the physiological situation with peptide-MHC, which do not all bind and dissociate at the same time.

Our optogenetic system allowed us to show that T cells discriminate between ligands due to differences in the ligand-TCR half-lives (Fig. 4D), consistent with KPR models (*6–8, 11*). Using the identical ligand-TCR pair for the different half-lives excludes potential differences in binding geometry (*12*), forces (*14, 15*) or conformational changes as discriminatory parameters in this setup. Furthermore, our results did not depend on the correlation of affinity or rate measurements from surface plasmon resonance experiments. On the contrary, we determined the binding parameters in the same experimental setup as the calcium stimulation experiments. The KPR duration of 16 s is well within the range of published binding half-lives of altered peptide ligands (*9, 28*). Future studies could investigate different factors impacting on the KPR duration, such as varying expression or activity levels of the involved kinases and phosphatases. Interestingly, changing the half-life of PhyB ON and thus the lifetime of the ligand-TCR interaction also alters the amount of bound receptors, which has broader implications for optogenetics.

In accordance with our results using soluble TCR ligands and the complete TCR complex, the study by Tischer et al. in this issue presents highly similar conclusions employing immobilized ligands and a chimeric antigen receptor.

Our approach could be a blueprint to study other ligand-receptor pairs and to understand how the kinetics of protein-protein interactions governs the activity of these binding events in diverse biological systems.

## Acknowledgements

We thank Susana Minguet for discussions and reading of the manuscript, the BIOSS toolbox for the equipment (INST 39/899–1 FUGG) to conduct the calcium experiments and Opto Biolabs for their customized illumination devices. This work was funded by the Deutsche Forschungsgemeinschaft (DFG) through EXC294 (Center for Biological Signalling Studies, BIOSS, WWS), and GSC-4 (Spemann Graduate School, OSY and MH). TH is a member of CellNetworks.

## Supplementary Materials

Materials and Methods

Supplementary Text

Fig S1 – S13

Table S1

References (25 – 41)

